# PSSR2: a user-friendly Python package for democratizing deep learning-based point-scanning super-resolution microscopy

**DOI:** 10.1101/2024.06.16.599221

**Authors:** Hayden Stites, Uri Manor

## Abstract

PSSR2 improves and expands on the previously established PSSR (Point-Scanning Super-Resolution) workflow for simultaneous super-resolution and denoising of undersampled microscopy data. PSSR2 is designed to put state-of-the-art technology into the hands of the general microscopy and biology research community, enabling user-friendly implementation of PSSR workflows with little to no programming experience required, especially through its integrated CLI and Napari plugin.

## 1 Introduction

The acquisition of high-quality microscopy data is time consuming and imperfect. Improvements to one of image resolution, SNR (Signal to Noise Ratio), imaging speed, or sample preservation cannot come without cost to another, requiring difficult and undesirable tradeoffs for imaging experiments. To mitigate this effect, PSSR (Point-Scanning Super-Resolution) was introduced to optimize these parameters to otherwise unattainable quality [1]. Using deep learning-based super-resolution, low quality images (low resolution, low SNR) could be restored to higher quality (high resolution, high SNR). Additionally, the use of a “crappifier” to semi-synthetically degrade high quality images to their low quality counterparts via downsampling and noise injection removed the requirement of a training dataset comprising perfectly aligned images, allowing for model training utilizing only high-resolution ground truth images.

However, the original version of PSSR was limited in implementation. For example, it only allowed for super-resolution of simple single-frame and multi-frame videomicroscopy timelapse data, not accounting for the usage of more complex multidimensional data where the benefits of PSSR may be even greater. The crappifiers used to generate training data were also imperfect, potentially contributing to decreased model accuracy when applied to real-world data.

Additionally, PSSR was also not designed with the bulk of the biological and imaging research community in mind. As machine learning becomes more widely established, the accessibility of related software becomes even more crucial. An experimental microscopist does not necessarily have experience in machine learning and/or any given programming language, and may not be able to fully benefit from, if at all, a workflow they cannot easily utilize. Underscoring this, the codebase was not structured as software, rather as a collection of undocumented functions that are not sufficient for customized usage without additional implementation. Additionally, the original code no longer runs on modern versions of the packages it is built on, which has caused difficulty for many prospective users. Thus, even though PSSR was released as open-source, it failed to maximize democratization.

To extend the democratization of PSSR, we introduce PSSR2, a Python package featuring a customizable framework for the creation of deep learning-based super-resolution/denoising imaging workflows on a wide variety of microscopy data including both light and electron microscopy. To this end, we redesigned the PSSR approach from the ground up with ease of use and wide applicability in highest priority, while also addressing the longstanding issues facing the original implementation.

The usage of the Python programming language allows for a user-friendly modular package-based interface and integration with widely used research software, such as PyTorch [2] and Napari [3]. PSSR2 is accessible through multiple entry points of varying complexities and required technical knowledge, while all including the same core features. For advanced users, lower level features are exposed in a way that allows for straightforward modification of existing processes and addition of custom logic. PSSR2 also boasts newly trained models with state-of-the-art performance on real-world data, substantially improving on prior PSSR implementations and other competing approaches.

## 2 Application Features

In order to produce a straightforward yet widely applicable PSSR2, we completely rebuilt PSSR from the ground up. In doing so, we simplify and improve important elements of the PSSR workflow while simultaneously allowing PSSR2 to properly run on a wider range of environments and dependency versions.

The PSSR2 package comprises six submodules under the main `pssr` module.

- `train`: Training and optimization functions
- `predict`: Prediction and benchmarking functions
- `data`: Datasets and functions for handling and generating paired high-low-resolution image pairs
- `crappifiers`: Pre-implemented and user definable crappifiers for semi-synthetically degrading high-resolution images into their low-resolution counterparts
- `models`: Pre-implemented neural network architectures
- `util`: Various utility functions

The majority of workflow configuration can be done solely in the selection and definition of objects within these modules. For many use cases, all relevant logic is predefined within corresponding objects, allowing users to remove unnecessary boilerplate code if desired. If accessing PSSR2 via its package interface, the user is not required to write more than a few lines of code defining objects to train and/or run a state-of-the-art model.

Perhaps the greatest source of workflow variance is in the input format and application of microscopy data. To efficiently provide a standardized interface for data loading, we separate our datasets into two categories: image datasets, which deal with preprocessed images, and sliding datasets, which deal with unprocessed data across multiple tiles within a larger image (i.e. a large image to be divided into multiple smaller images). Tiles are defined by horizontal and vertical positions within an image if applicable, where any additional dimensional data is separated into multiple frames of the same tile as set by a user-defined configuration. We separate training and validation sets randomly by tile, rather than by frame, because different frames from the same tile may be overly similar to one another, which may lead to undesirable training data leakage or overfitting. Both dataset types are capable of model training and prediction. In training, datasets automatically generate semi-synthetic low-resolution images from given high-resolution images using a configurable “crappifier”. For model predictions, datasets load only low-resolution images to be restored (i.e. “decrappification”). For model benchmarking, we also provide datasets that omit the crappification of high-resolution images, instead loading real-world high-low-resolution image pairs.

We designed datasets in this manner to be effectively “plug-and-play” while still being comprehensive for as many applications and data types as possible. Additionally, the usage of n-dimensional images is supported, including those with asymmetrical amounts of dimensions between model inputs and outputs, along with the training of models for image denoising tasks. While models could be trained on multidimensional images in the prior PSSR implementation, we expand the scalability and viability of such an approach beyond the single-frame and multi-frame temporal super-resolution approaches previously used.

Various pre-implemented neural network architectures for image super-resolution are accessible from within the package. The included implementations are modified to strip away excess dependencies and bloat, and are standardized to be easily understandable for inexperienced users. However, these are solely for ease of use, and more advanced users are free to use any PyTorch-based neural network architecture.

As the majority of PSSR2 functionality was designed to be accessed through simple object interfaces, it allows for the seamless implementation of said functionality into external applications. In addition to accessing PSSR2 as a Python package, it is also available as a Command Line Interface (CLI) and a Napari graphical user interface plugin, each with the same core features as the Python package. Both the CLI and Napari plugin allow training and prediction using custom defined datasets and models. Additionally, the latter allows for runtime monitoring of model output images in multidimensional spaces within the Napari viewer.

All functions and objects are fully documented in docstrings visible in interactive programming environments, which are also available along with a user guide in the online documentation.

## 3 Advancements in PSSR2

PSSR2 overhauls the semi-synthetic data generation process (i.e. “crappification”) of its predecessor, contributing to increased robustness and prediction accuracy on real-world data. In the crappification process of the prior PSSR implementation, semi-synthetic low-quality images (low resolution, high noise) are generated from real-world high-quality images (high resolution, low noise) by adding noise and then downsampling. However, by adding noise first, adjacent noise values are averaged together in downsampling contributing to lower noise variance, especially in value dependent noise distributions such as Poisson noise. Though a seemingly minor change, we found that by adding noise after downsampling, semi-synthetically generated low-resolution training images are more representative of real-world images, thereby enabling more accurate model training. We found that this allowed Poisson noise to be used to great success, where even though it is most statistically similar to the real-world noise processes in a point-scanning system, the prior implementation found it performing poorly.

We also allow training set images to be crappified on statistically independent noise intensity values rather than a set intensity across all images, which allows trained models to function on a wider range of input data qualities. Additionally, by generating training images during runtime rather than before training as did the previous implementation, it allows the same image to receive many distinctly different crappifications (random noise, random intensity) across all training epochs. With both changes in data augmentation combined we can effectively increase the size of the training dataset by a large factor, reducing overfitting and increasing robustness.

We also developed a methodology for measuring the amount of noise present in a low-resolution image when compared to its corresponding high-resolution image for use in properly quantifying crappifier intensities for more accurate training data generation. The high-resolution image is first downsampled to create a “noiseless” low-resolution image, the values of which are then subtracted from the true low-resolution image to yield a noise profile, i.e. an n-dimensional array representing the noise in the low-resolution image. Crappifier parameters are then approximated by Bayesian optimization using Gaussian processes [4], minimizing the distribution of noise profile values of the semi-synthetic low-resolution image generated from fit crappifier parameters against that of the true low-resolution image.

We benchmarked the real-world improvements of PSSR2 using a test dataset consisting of 42 pairs of both high and low resolution images. Performance is measured as the ability of the model to restore the low-resolution images to their high-resolution counterparts, with a simple bilinear upscaling operation used as a control. In our testing, we found that PSSR2 increased model prediction accuracy in terms of both PSNR (Peak Signal-to-Noise Ratio) and SSIM (Structural Similarity) metrics when compared to the prior PSSR implementation (Fig. 1b). Strikingly, the SSIM performance uplift of PSSR2 against the control approximately doubled that of the prior PSSR implementation. As SSIM measures higher order visual changes than PSNR, this can be attributed to greater visual clarity and more accurate texture in PSSR2 output.

**Figure 1:**
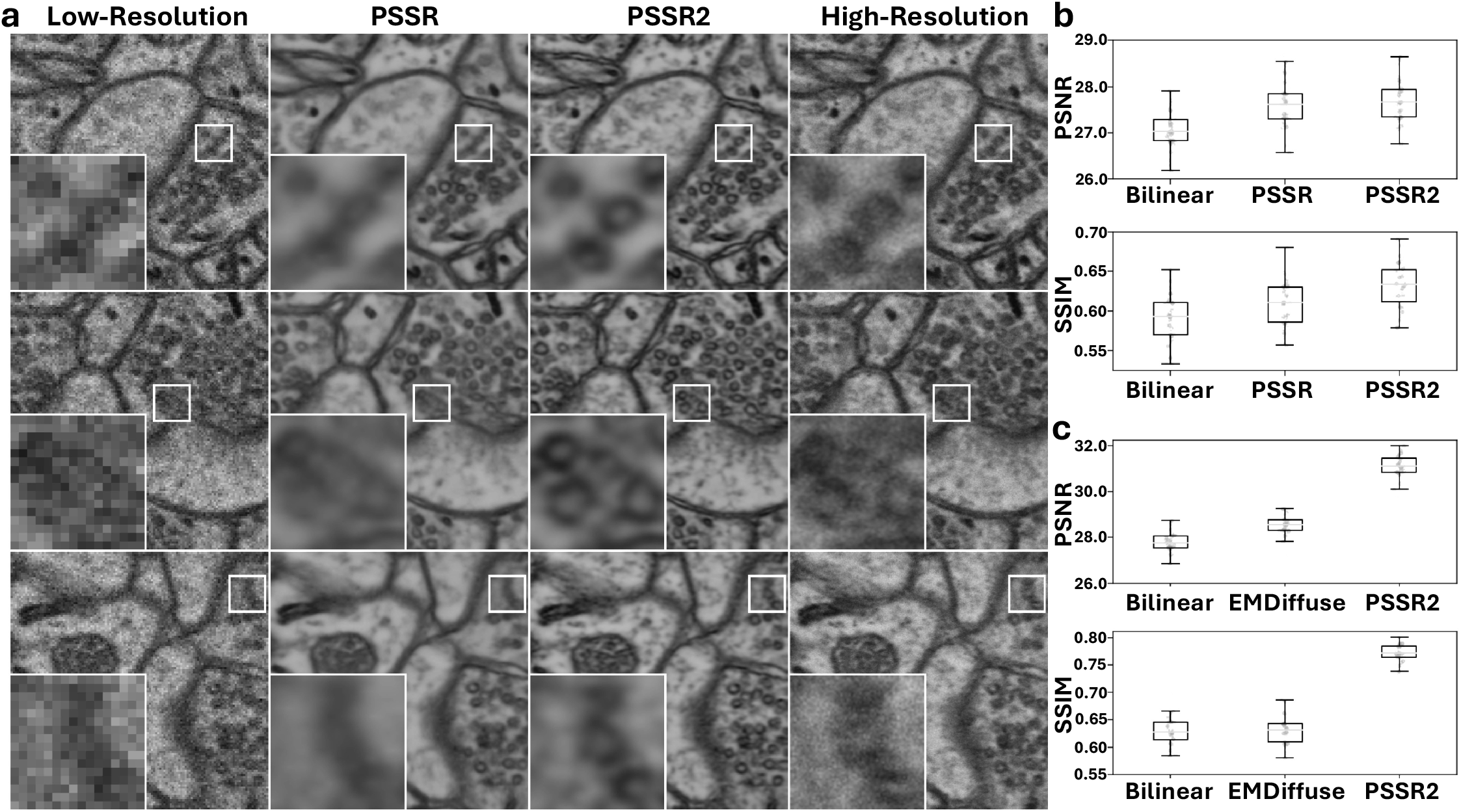
Evaluation of PSSR2 against state-of-the-art microscopy image restoration methods. a, Super-resolution ResUNet model predictions of both PSSR2 and the prior PSSR implementation, in comparison to low-resolution inputs and high-resolution ground truth. b, Image restoration performance comparison of ResUNet super-resolution models trained with either the prior PSSR implementation or PSSR2, with a bilinear upscaling control. c, Image denoising performance of PSSR2-trained ResUNet model against an EMDiffuse-trained model, with a bilinear upscaling control.

In visual examination of images generated by both PSSR2 and the prior PSSR implementation each using the same low-resolution model input images, the increase in image clarity in PSSR2 generated images is apparent, with less blur than its predecessor (Fig. 1a). The elimination of these artifacts, while maintaining robust model performance, should improve the performance of any additional or downstream algorithms/workflows.

To prove the merits of PSSR2 on multiple training tasks, we also benchmarked image denoising against EMDiffuse, another state-of-the-art microscopy image restoration tool utilizing diffusion-based deep learning models [5]. We found that the image denoising predictive accuracy of PSSR2 surpasses that of EMDiffuse in terms of both PSNR and SSIM metrics (Fig. 1c), proving its usefulness in a variety of scenarios. In visually examining image predictions from both algorithms, PSSR2 tends to only predict detail that can be reasonably inferred from the low-resolution input image, whereas EMDiffuse tends to infer beyond the data represented in the image. While this quality could be useful in applications with extreme amounts of image noise, especially because the architecture also models prediction uncertainty, it ultimately generates images that, while perhaps visually appealing, do not accurately represent the hidden information in the low-resolution inputs. This demonstrates what is perhaps the greatest strength of PSSR2, in that it is able to restore a detailed image while maintaining a low rate of false positive image elements, also known as hallucinations.

For both use-cases, our benchmarked models were trained with MS-SSIM + L1 loss [6]. We found this loss function provided faster model convergence and improved visual clarity over L2 loss, which was used to train the model in the prior PSSR implementation.

## 4 Conclusion

We present here a user-friendly and robust Python package capable of state-of-the-art deep learning-based microscopy super-resolution, easily accessible through its integrated CLI and Napari plugin in addition to a Python interpreter. Offering many configurable objects and tools for a variety of use cases while being concise and approachable, PSSR2 is designed to be accessible to most imaging and biology researchers through its simple yet expandable interface. PSSR2 utilizes the widely used PyTorch library, allowing for simple integration with state-of-the-art models and architectures. Importantly, PSSR2 expands on the prior implementation of PSSR to provide more realistic and efficient model training on a wide variety of microscopy data and use-cases. PSSR2 boasts a significant increase in image quality from its predecessor, and outperforms other image restoration approaches in accuracy. We hope this update provides a meaningful step towards democratizing access to image enhancement tools for the biological imaging community.

## Acknowledgments

U.M. is supported by the Goeddel Family Technology Sandbox at the UC San Diego, NIDCD R01 DC021075, an NSF NeuroNex Award (2014862), and the CZI Imaging Scientist Award DOI https://doi.org/10.37921/694870itnyzk from the Chan Zuckerberg Initiative DAF, an advised fund of Silicon Valley Community Foundation (funder DOI https://doi.org/10.13039/100014989).

We would like to thank Linjing Fang for the original implementation of PSSR and assistance in accessing the training/testing data, and Yasmin Kassim for helpful insights and feedback.

## Availability and Implementation

PSSR2 is a cross-platform Python package distributed on the Python Package Index. The source code for the CLI and Napari plugin are available under the MIT license at https://github.com/ucsdmanorlab/PSSR2.

## Supplementary Information

The PSSR2 documentation, containing a comprehensive user guide, full API reference, and links to pretrained models and sample datasets are available at https://ucsdmanorlab.github.io/PSSR2.

